# Effects of 6-methyl nicotine in middle aged female rats with a history of nicotine vapor self-administration

**DOI:** 10.1101/2025.09.15.676338

**Authors:** Michael A. Taffe, Tyra R. Coons, Tess A. Doran, Yanabel Grant, Sophia A. Vandewater

## Abstract

**Rationale:** The nicotine analog 6-methyl nicotine (6-MN) has recently appeared in non-tobacco nicotine delivery products, including oral pouches and e-cigarette liquids, in an apparent ploy to evade regulation of nicotine by the United States Food and Drug Administration or other public health agencies. Unfortunately, only minimal scientific information on the effects of 6-MN is available.

**Objective:** To determine the extent to which 6-MN produces nicotine-like effects on body temperature, wheel activity and nociception in laboratory rodents.

**Methods:** Middle-aged (starting at Post Natal Day 425) female Wistar rats were evaluated for rectal temperature, voluntary wheel activity, and nociceptive responses (warm water tail-withdrawal) to subcutaneous injection of nicotine (0.0, 0.8 mg/kg) or 6-methyl nicotine (6-MN; 0.0, 0.4, 0.8 mg/kg). Temperature and nociceptive responses to vapor inhalation of 6-MN [0-30 mg/mL in the propylene glycol (PG) vehicle] were assessed. Finally, the self-administration of 6-MN vapor was compared with nicotine vapor self-administration.

**Results:** 6-MN decreased rectal temperature, suppressed wheel activity and induced modest nociceptive effects. The magnitude of the effect of 0.8 mg/kg 6-MN and 0.8 mg/kg nicotine were similar across all three assays. Vapor self-administration of 6-MN and nicotine was likewise comparable at a 10 mg/mL concentration.

**Conclusion:** 6-MN administered by injection or by vapor inhalation produces behavioral and physiological effects that are very similar to those produced by nicotine in rats. It is therefore likely that detrimental health effects of 6-MN will be quite similar to those established for nicotine.

## 1. Introduction

Vapor inhalation of nicotine using a variety of electronic nicotine delivery systems (ENDS; “e-cigarettes”) exceeds the use of cigarettes for nicotine ingestion in adolescent and young adult populations by a considerable margin. Survey data from the United States show that in 2023 about 19% of 19-30 year olds had vaped nicotine in the past month whereas only 8.8% had used cigarettes (Patrick et al., 2024). Similarly, 16.9% of 12-grade students had vaped nicotine in the past 30 days, whereas only 2.9% used cigarettes (Miech et al., 2025). *Daily* use incidence for 12^th^ graders was 5.8% for vaping and 0.7% for cigarettes in 2023, and 5.2%/0.4% in 2024 (Miech et al., 2025); also see (Park-Lee et al., 2022).

Efforts by the United States Food and Drug Administration to regulate tobacco and nicotine products has led to an attempt by the retail industry to evade regulation by using 6-methyl nicotine (6-MN) as the active constituent (Jordt et al., 2024). Although 6-MN occurs in tobacco products, its abundance is ∼4 orders of magnitude less than nicotine (Pankow et al., 2025), and thus its effects have not been widely studied. 6-MN has been recently confirmed in oral pouch products (Jordt and Jabba, 2024; Mallock et al., 2024; Vanhee et al., 2024) and in ENDS liquids (Jordt et al., 2024) as a primary psychoactive constituent. A recent survey found that ∼20% of adolescents (oversampled for prior tobacco use) were aware of, and 8% had used, a specific nicotine analogue product that had entered the market six months prior (Sanchez et al., 2025). The website for one commercial disposable pod e-cigarette product (SpreeBar, 2023) claims their active substance to be “*a patented non-nicotine compound*” and “*A synthetically derived molecule that is structurally similar to, but chemically different from other vaping alkaloids, Metatine is not made from, and does not contain, nicotine*” and this product has been shown by chemical analysis to contain 6-MN (Jordt et al., 2024). One study reported a content of ∼6 mg of 6-MN per gram of e-liquid in several products that were labeled as containing 6-MN at 5%, or 50 mg/g (Erythropel et al., 2024), suggesting the producers of these products think 6-MN is ∼8-fold more potent than nicotine. While one study found that (-)-6-MN was about 3-fold more potent than (-)-nicotine in a competition binding assay in rat membrane (Wang et al., 1998), another indicated that (+/-)-6-MN was about 3-fold more potent than (+/-)-nicotine in tail-withdrawal anti-nociception and 5-fold more potent in inhibiting spontaneous activity in mice (Dukat et al., 2002). A report from within the tobacco industry found 6-MN equipotent with nicotine in a drug-discrimination assay in male rats (Levy and Dunn, 1979). This disagreement of evidence on potency underlines the need for additional laboratory study.

Controlled animal models are necessary to more precisely delineate risks associated with ENDS based inhalation of nicotine and analogue replacements such as 6-methyl nicotine, given social factors which influence teen smoking and vaping experiences. There is at present very little scientific information on 6-MN available, indeed, a recent scoping review of toxicity associated with 6-MN ended up being mostly speculation drawn from indirect evidence (Effah et al., 2025). Recent work shows that ENDS devices can deliver physiologically effective doses of nicotine to laboratory rat subjects by inhalation, altering body temperature, spontaneous locomotor activity and nociception (Javadi-Paydar et al., 2019; Javadi-Paydar et al., 2024; Montanari et al., 2020). Additional work from several labs shows that nicotine vapor delivery reinforces operant responding in rats (Gutierrez et al., 2024a; Gutierrez et al., 2024b; Lallai et al., 2021; Smith et al., 2020); self-administration has also been demonstrated in mice (Cooper et al., 2021; Henderson and Cooper, 2021). Thus, rat models can be deployed to determine the extent to which 6-MN has behavioral pharmacological properties similar to, or different from, those of nicotine.

This study was designed to examine the impact of 6-MN in a model of long-term nicotine exposure in middle aged female rats. Epidemiological data show an increase in 30 day prevalence of smoking from 8.5% to 11.2% of women age 55-65 from 2023 to 2024 and about 8-9% incidence of daily cigarette use from age 35 to 65 (Patrick et al., 2025). Early adult female rats were exposed from Post-Natal Day 115 onward to twice-daily inhalation of nicotine vapor (30 or 60 mg/mL in the Propylene Glycol vehicle) or that of the vehicle over 10 days and assessed from PND150 to PND 390 (∼5 to 13 months of age) for nicotine vapor self-administration in 81 sessions. Female laboratory rats show alterations of neuroendocrine markers that are consistent with middle age (Downs and Wise, 2009), start to develop irregular estrus cycles around 12 months of age (Lu et al., 1994) and rats are studied as cognitively aged around 21 months (Gonzalez-Hedstrom et al., 2021; Rojic-Becker et al., 2021) with 24 months representing an effective group lifespan in laboratory studies. These rats were therefore selected as a model of a lifetime nicotine consumer who encounters one of these novel 6-MN products. The study first compared the effects of 6-MN to nicotine after subcutaneous injection on measures of body temperature, anti-nociception, and voluntary wheel activity. Experiments next assessed the impact of 6-MN when delivered by vapor inhalation using an e-cigarette style device. Finally, studies evaluated the propensity for vapor self-administration of 6-MN compared with the vapor self-administration of nicotine.

## 2. Methods

### 2.1 Subjects

Female (N=36) Wistar rats obtained from the vendor (Charles River) on Post Natal Day (PND) 74 were used for this study. The vivarium was kept on a 12:12 hour reversed light-dark cycle, and behavior studies were conducted during the vivarium dark period. Food and water were provided *ad libitum* in the home cage. Procedures were conducted in accordance with protocols approved by the Institutional Animal Care and Use Committee of the University of California, San Diego and were consistent with the NIH Guide (Garber et al., 2011). The rats had been exposed to 30 minute vapor inhalation sessions, twice daily at ∼5 h intervals, from PND 115-119 and PND 122-126 with N=3 per chamber. Groups (N=12) were exposed to vapor from the propylene glycol (PG) vehicle or with 30 or 60 mg/mL (-)-nicotine bitartrate in the PG. All groups self-administered (-)-nicotine vapor in 81 sessions conducted from PND-150 to PND-390 under various concentration (30 mg/mL training dose; 3-60 mg/mL in dose-substitution) and pre-treatment with synthetic cooling agents (0.3-30 mg/kg, N-Ethyl-p-methane-3-carboxamide; WS-3, and 2-isopropyl-n,2,3-trimethylbutanamide; WS-23) conditions. No major differences in self-administration behavior associated with original exposure group were observed after the initial acquisition period.

### 2.2 Drugs

(-)-Nicotine bitartrate (Sigma-Aldrich, St. Louis, MO) or (-)-6-methyl nicotine (Nicotine River; Thousand Oaks, CA; and NIDA Drug Supply Program) were dissolved in propylene glycol (PG; Fisher Scientific) at concentrations of 5, 10 and 30 mg/mL for vapor studies. (-)-PG was used as the vehicle for consistency and comparability with our prior reports on the impact of repeated nicotine vapor inhalation (Gutierrez et al., 2024a; Gutierrez et al., 2024b). Nicotine bitartrate (Sigma-Aldrich, St. Louis, MO), or (+/-)-6-methyl nicotine bitartrate (NIDA Drug Supply Program) were dissolved in physiological saline for subcutaneous injection studies.

### 2.3 Apparatus

An Electronic Nicotine Delivery System (ENDS) based vapor inhalation apparatus (La Jolla Alcohol Research, Inc) was used for these studies. In brief, vapor was delivered into sealed vapor exposure chambers through the use of controllers which trigger commercial e-cigarette tanks/atomizers (SMOK TFV8 X-baby; 0.25 ohm V8 X-Baby M2 Core). Vacuum control through an exhaust valve flowed room air at ∼1 L per minute and ensured that vapor entered the chamber on each device triggering event. Puffs were delivered every 5 minutes during the session for non-contingent exposure.

### 2.4 Experiments

#### 2.4.1 Experiment 1: Effect of nicotine and 6-methyl nicotine on temperature

Administration of (-)-nicotine ditartrate by s.c. injection or vapor inhalation reduces the core body temperature of rats (Javadi-Paydar et al., 2018; Javadi-Paydar et al., 2019). A group of rats (N=12; N=4 from each of the repeated PG, Nicotine 30 and Nicotine 60 groups) were used for this experiment. Studies were conducted from PND-425 to PND-445 with a minimum of three days between evaluations in any given rat. For this study, a baseline temperature was obtained prior to injection and then re-assessed at 30, 60 and 120 minutes after injection. The impact of (-)-nicotine (0.0, 0.8 mg/kg, s.c.) was first assessed in a balanced order, and then the impact of (+/-)-6-MN (0.0, 0.4, 0.8 mg/kg, s.c.), were next assessed in a balanced order. Injection of one rat (0.4 mg/kg (+/-)-6-MN) was not successful thus this point was excluded from the analysis.

Four weeks later, rectal temperature was assessed before and 35 and 60 minutes after the initiation of a 30 minute vapor inhalation session. Conditions were counterbalanced to assess the effect of concentrations of (-)-6-MN (0, 5, 10 and 30 mg/mL in the PG vehicle). Studies were conducted from PND-473 to PND-487 with a minimum of three days between evaluations in any given rat.

#### 2.4.2 Experiment 2: Effect of nicotine and 6-methyl nicotine on wheel activity

Administration of (-)-nicotine ditartrate reduces voluntary wheel activity in rats when administered by s.c. injection or vapor inhalation (Gutierrez et al., 2024a; Gutierrez et al., 2024b). A different group of rats (N=8; N=2 repeated PG group; N=3 repeated Nicotine 30 and Nicotine 60 groups) were used for this experiment. Studies were conducted from PND-425 to PND-445 with a minimum of three days between evaluations in any given rat. Rats were injected 15 minutes prior to being provided wheel access, with activity being measured in quarter-rotations every 5 minutes. Injection of one rat for the 0.8 mg/kg nicotine session was not successful, thus this animal was excluded from that analysis.

#### 2.4.3 Experiment 3: Effect of nicotine and 6-methyl nicotine on nociception

A third subset (N=8; N=3 from the repeated PG and repeated Nicotine 30 groups; N=2 from the Nicotine 60 group) were evaluated on nociceptive responses using a warm water (52 °C) tail-withdrawal assay. The impact of vehicle (saline), (-)-nicotine (0.8 mg/kg, s.c.) and (+/-)-6-MN (0.4, 0.8 mg/kg, s.c.) were evaluated, with all four conditions assessed in a counter-balanced order. A pre-injection baseline withdrawal latency was obtained and then the assay repeated 15, 30 and 60 minutes after injection. Studies were conducted from PND-453 to PND-465 with a minimum of three days between evaluations in any given rat.

The rats were next assessed after vapor inhalation of PG, 30 mg/mL (-)-nicotine or 30 mg/mL (-)-6-MN in a counter-balanced order. Studies were conducted from PND-480 to PND-486 with a minimum of three days between evaluations in any given rat.

#### 2.4.4 Experiment 4: Effect of 6-methyl nicotine vapor inhalation on nociception and body temperature

A fourth subset (N=8; N=2 repeated PG, and N=3 repeated Nicotine 30 and Nicotine 60 groups) were evaluated on nociceptive responses using a warm water (52 °C) tail-withdrawal assay and thermoregulation using a rectal thermister. These individuals were PND 495 at the start of this study and had not been exposed to any nicotine since their final nicotine vapor self-administration session on PND 390. Tail withdrawal and rectal temperature were assessed prior to a 30 minute inhalation session, 35 minutes after the start of inhalation and 60 minutes after the start of inhalation. Inhalation conditions of PG, and (-)-6-MN (5, 10, 30 mg/mL in the PG vehicle) were assessed in a counterbalanced order with at least 3 days between assessments.

#### 2.4.5 Experiment 5: Self-administration of 6-methyl nicotine by vapor inhalation

A group of the middle aged female rats used for the above experiments (N=20; N=7 repeated PG and repeated Nicotine 30 groups and N=6 repeated Nicotine 60 group) were evaluated on inhalation self-administration after the above experiments, beginning on PND 493. This was approximately 15 weeks after their prior vapor self-administration session, which was used to assign the groups based on ∼equivalent mean deliveries obtained. For this study, puffs of vapor from either (-)-6-MN (N=10; 10 mg/mL concentration) or (-)-nicotine (N=10; 10 mg/mL concentration) were made available upon a lever response under a Fixed Ratio 1 contingency in 30-minute sessions. After seven sessions, the groups were permitted to respond for vapor with the alternate drug for another seven sessions.

### 2.5 Data Analysis

Rectal temperature (°Celsius), wheel quarter-rotations and tail withdrawal latency (seconds) were analyzed by 2-way ANOVA including within-subjects factors for Dose condition and Time. One-way ANOVA was used with a within-subject factor of Dose condition to compare wheel activity session sums, the 30 minute post-injection timepoint for rectal temperature and the 15 minute post-injection timepoint for tail withdrawal latency. The self-administration data (vape deliveries and percent of responses on the drug-associated lever) were analyzed by group and by session, within each set of 7 sessions conducted for a single drug assignment. Additional analysis compared the 7-session average for each drug condition (6-MN vs nicotine) within and across groups. Mixed effects analysis was used in any cases where data points were missing.

In all analyses, a criterion of P<0.05 was used to infer a significant difference. Significant main effects were followed with post-hoc analysis using Tukey (multi-level factors), Sidak (two-level factors) or Dunnett (treatments versus control within group) correction. All analysis used Prism for Windows (v. 10.5.0; GraphPad Software, Inc, San Diego CA).

## 3. Results

### 3.1 Experiment 1: Effect of nicotine and 6-methyl nicotine on temperature

Nicotine injection decreased temperature (**Figure 1A**) as confirmed by significant effects of Time [F (1.284, 14.12) = 6.59; P<0.05], Dose [F (1.000, 11.00) = 32.03; P<0.0005] and of the interaction of Time with Dose [F (1.602, 17.62) = 8.26; P<0.005] in the ANOVA. The Tukey post-hoc test confirmed significant differences in temperature compared with saline 30-120 minutes after injection. Likewise, body temperature 30 minutes after nicotine injection differed significantly from the pre-injection and 60-120 minute post-injection timepoints; the latter also differed significantly from each other.

Injection of 6-MN had similar hypothermic effect (**Figure 1B**) as confirmed by significant effects of Time [F (1.242, 13.66) = 37.41; P<0.0001], Dose [F (1.811, 19.92) =10.56; P<0.005] and of the interaction of Time with Dose [F (3.266, 33.75) = 14.79; P<0.0001] in the mixed-effects analysis (one individual had a missing value for 0.4 mg/kg, due to incomplete injection). The Tukey post-hoc test confirmed significant differences in temperature between all conditions 30 minutes after injection and between the 0.8 mg/kg 6-MN and saline conditions 60 minutes after injection. There was also a significant difference in temperature 30 minutes after 0.4 or 0.8 mg/kg compared to all other timepoints within-condition and 60-120 minutes after 0.8 mg/kg compared with the pre-injection temperature.

**Figure 1:**
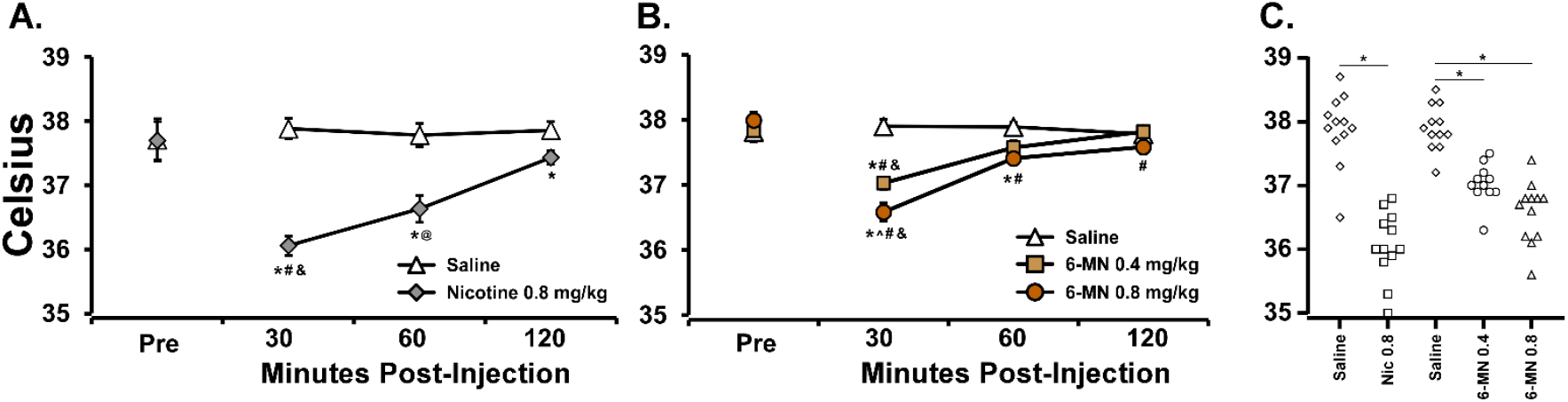
Mean (N=12; ±SEM) rectal temperature before and after injection with **A)** Nicotine (0.0, 0.8 mg/kg, s.c.) or **B)** 6-methyl nicotine (6-MN; 0.0, 0.4, 0.8 mg/kg, s.c.), and **C)** Individual 15-minute post-injection temperatures for all five treatment conditions. A significant difference from saline at the respective timepoint is indicated with *, from the 0.4 mg/kg dose with ^, from the pre-injection baseline with #, from both other post-injection times with & and from the 120 minute timepoint with @.

One way mixed effects analysis of the 30 minute post-injection temperatures including all five treatment conditions confirmed a significant effect of treatment condition [F (2.454, 33.13) = 36.69, P<0.0001]. The Tukey post-hoc test confirmed a significant difference from the respective saline conditions after 0.8 mg/kg nicotine and both doses of 6-MN, but did not confirm a significant difference between nicotine 0.8 mg/kg and 6-MN 0.8 mg/kg (**Figure 1C**).

### 3.2 Experiment 2: Effect of nicotine and 6-methyl nicotine on wheel activity

Nicotine injection reduced wheel activity (**Figure 2A**), as the ANOVA confirmed that wheel activity was significantly affected by Time [F (2.842, 17.05) = 12.05; P<0.0005], by Drug Treatment Condition [F (1.000, 6.000) = 45.63; P<0.001] and by the interaction of Time with Drug Treatment Condition [F (2.734, 16.40) = 14.73; P<0.0001]. The post-hoc test limited to orthogonal comparisons confirmed that activity differed from saline at all timepoints except 20 minutes after injection. Similarly, 6-MN injection also reduced wheel activity (**Figure 2B**), as the ANOVA confirmed that wheel activity was significantly affected by Drug Treatment Condition [F (1.000, 6.000) = 45.63; P<0.001] and by the interaction of Time with Drug Treatment Condition [F (2.734, 16.40) = 14.73; P<0.0001]. The post-hoc test limited to orthogonal comparisons confirmed that activity differed from saline 5-10 minutes after the 0.4 mg/kg dose and 5-20 minutes after injection with 0.8 mg/kg. One-way mixed-effects analysis of the wheel activity session totals (**Figure 2C**) confirmed a significant effect of treatment condition [F (2.245, 14.59) = 18.52; P<0.0001]; the Tukey post-hoc test confirmed that wheel activity was lower after the 0.8 mg/kg dose of either nicotine or 6-MN compared with either saline treatment.

**Figure 2:**
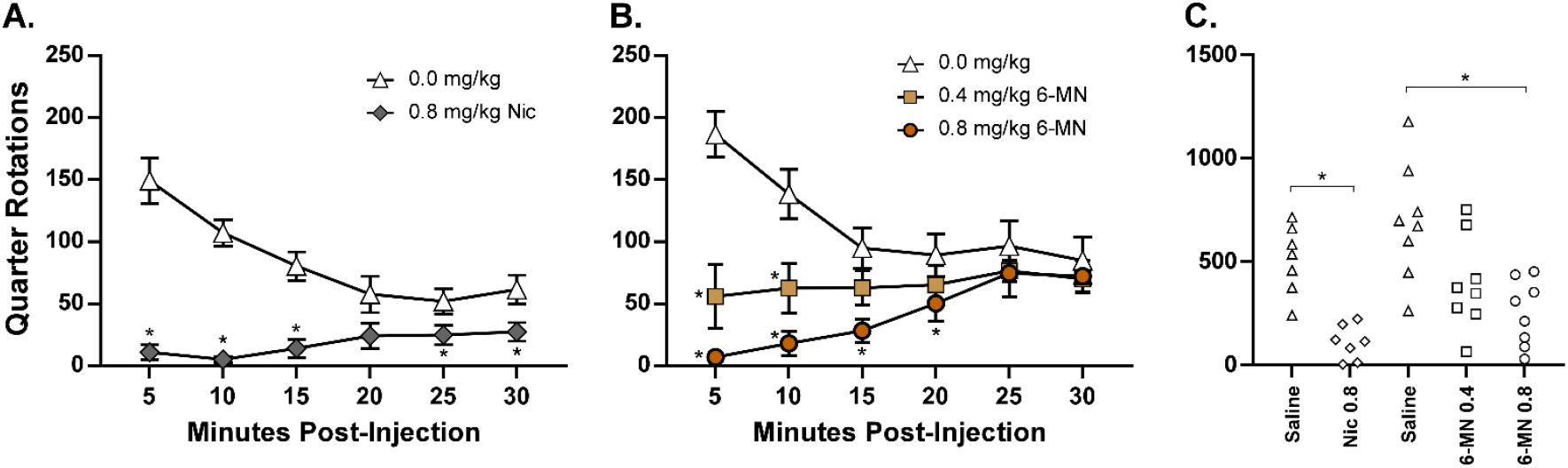
Mean (N=7-8; ±SEM) wheel activity after injection with **A)** Nicotine (0.0, 0.8 mg/kg, s.c.) or **B)** 6-methyl nicotine (6-MN; 0.0, 0.4, 0.8 mg/kg, s.c.), and **C)** Individual session wheel activity sum for all five treatment conditions. A significant difference from saline injection is indicated with *.

### 3.3 Experiment 3: Effect of nicotine and 6-methyl nicotine on nociception

The ANOVA confirmed that tail withdrawal latencies (**Figure 3**) were significantly affected by Time [F (3, 21) = 8.11; P<0.001] and by the interaction of Time with Drug Treatment Condition [F (9, 63) = 3.26; P=0.0026] but not by Time [F (3, 21) = 2.51; P=0.087]. The post-hoc test limited to orthogonal comparisons confirmed that latencies were significantly elevated 15 minutes after both 6-MN doses compared with saline and after 0.8 mg/kg 6-MN compared with 0.8 mg/kg nicotine. Within treatment condition, latencies were higher 15 minutes after 0.4 mg/kg 6-MN compared with the pre-injection and 60-minute post-injection values and higher 15 minutes after 0.8 mg/kg 6-MN compared with the pre-injection and the 30- and 60-minute post-injection values. The one-way ANOVA for the 15 minute latencies confirmed an effect of treatment condition [F (3, 21) = 4.0; P<0.05] and the Tukey post-hoc further confirmed significantly longer latency after the 0.8 mg/kg dose of 6-MN compared with saline.

**Figure 3:**
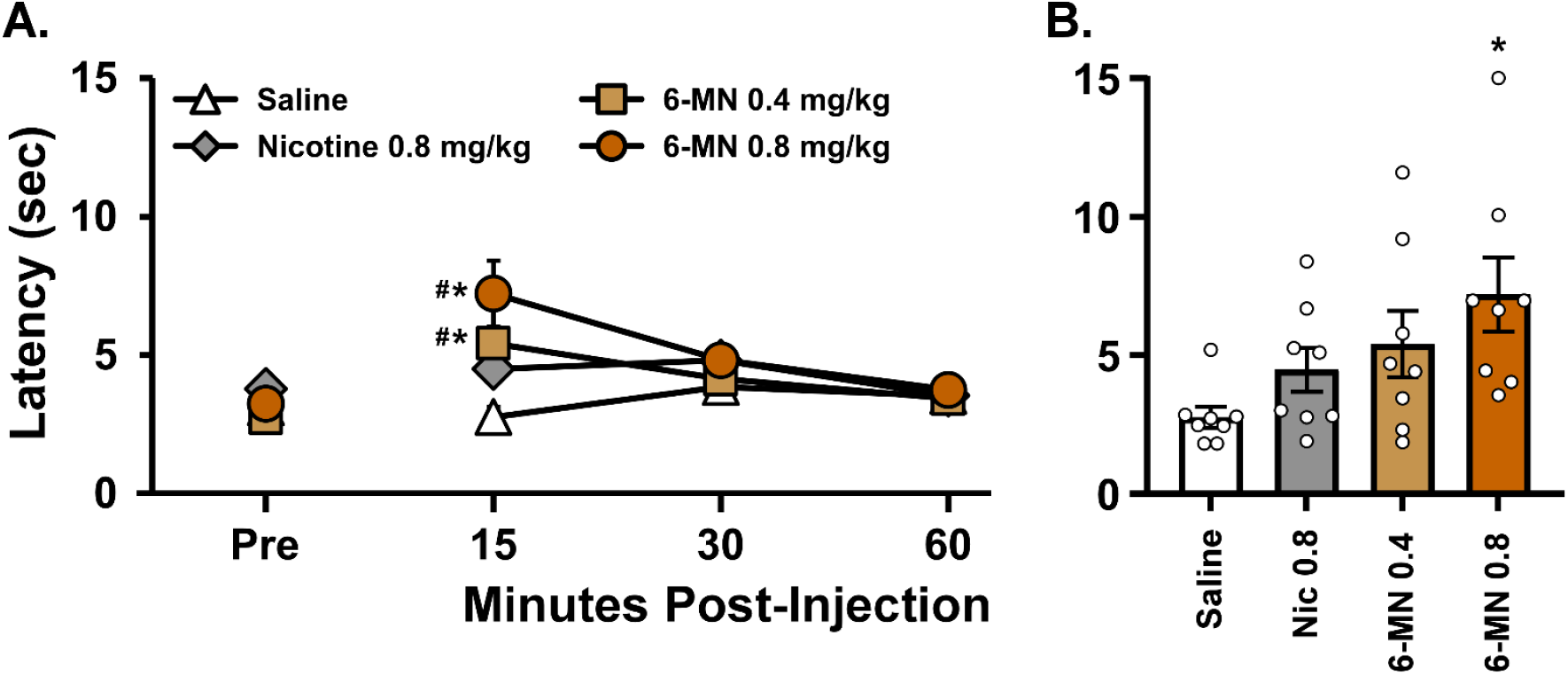
Mean (N=8; ±SEM) tail-withdrawal latency before and after injection with saline, nicotine (0.8 mg/kg, s.c.) or 6-methyl nicotine (6-MN; 0.4, 0.8 mg/kg, s.c.). A significant difference from saline is indicated with * and from the pre-injection baseline with #.

There was a minor significant effect of nicotine, but not 6-MN, confirmed in the vapor inhalation study (not shown). It is possible that some degree of tolerance was produced by the injection experiments in these animals, as we’ve reported before for animals in repeated longitudinal studies of nicotine (Javadi-Paydar et al., 2019) and as occurs in animals repeatedly treated with 1 mg/kg, s.c. (Epstein et al., 1989). This conclusion is supported by the successful alterations produced by the inhalation approach in a subset of animals who had not been exposed to nicotine or 6-MN for 105 days, see Experiment 4.

### 3.4 Experiment 4: Effect of 6-methyl nicotine vapor inhalation on nociception and body temperature

The inhalation of 6-MN decreased body temperature with a maximum mean effect of about 1.5 °Celsius (**Figure 4A**). The ANOVA confirmed that rectal temperatures were significantly affected by Time [F (1.826, 12.78) = 45.79; P<0.0001], by Drug Treatment Condition [F (2.388, 16.72) = 5.01; P<0.05] and by the interaction of Time with Drug Treatment Condition [F (2.323, 16.26) = 7.41; P<0.005]. The Tukey post-hoc test of all orthogonal comparisons confirmed temperature was lower 35 minutes after initiation of the 30 mg/mL condition compared with the PG and 5 mg/mL conditions. Similarly, temperature was significantly lower 35 minutes after the start of 10 mg/mL inhalation compared with the pre-treatment temperature and all three timepoints in the 30 mg/mL condition significantly differed from each other.

**Figure 4:**
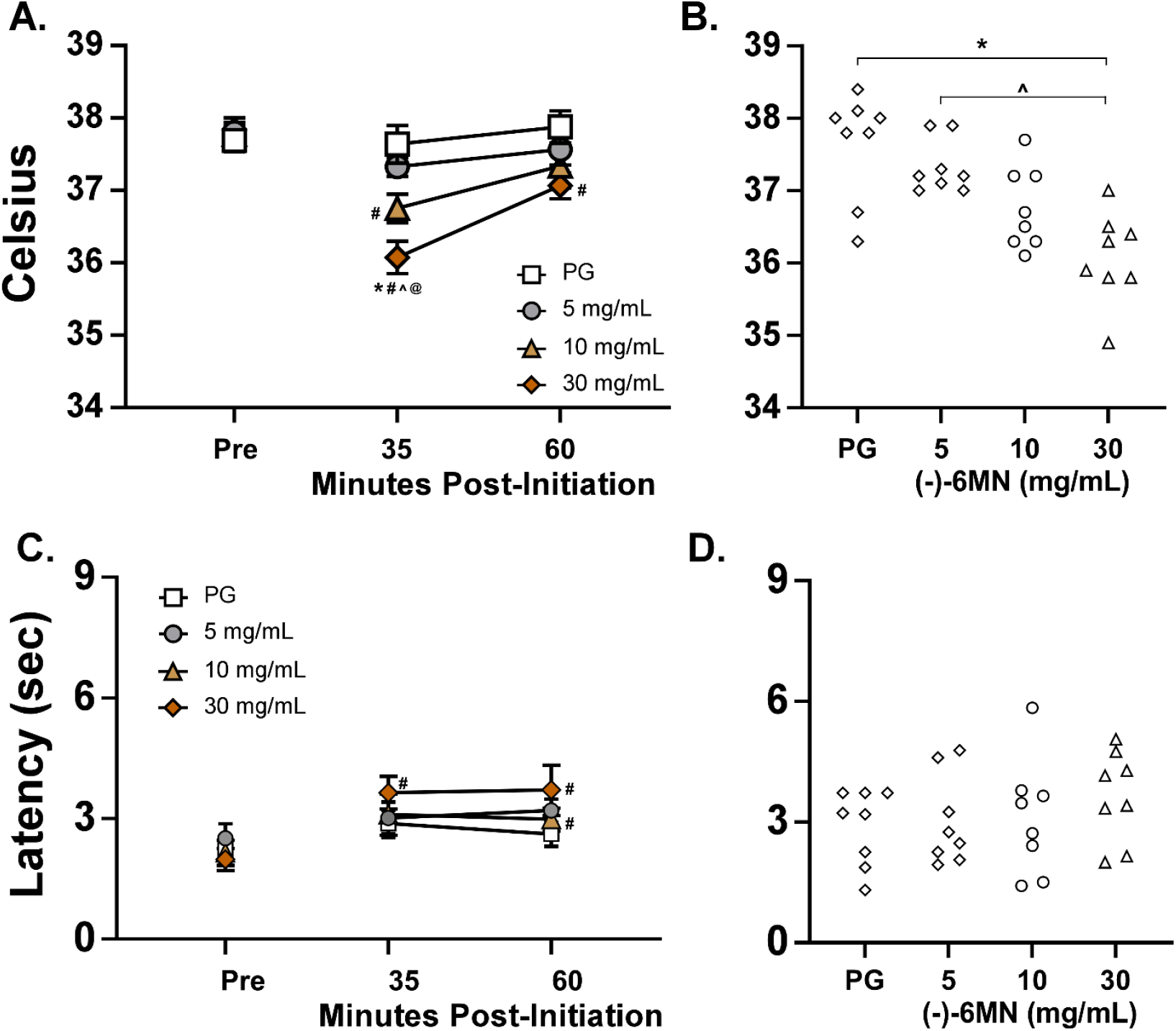
Mean (N=8; ±SEM) **A)** rectal temperature and **C)** tail-withdrawal latency assessed before and after inhalation of vapor from the PG vehicle or (-)-6-methyl nicotine (6-MN5, 10, 30 mg/mL in the PG). A comparison of individual **B)** temperatures and **D)** tail-withdrawal latencies observed 35-minute post-initiation is depicted for all four treatment conditions. A significant difference from vehicle vapor (PG) is indicated with *, from the 5 mg/mL concentration with ^, from the pre-inhalation baseline with # and from the 60 minute timepoint with @.

Nociception was also modestly decreased by 6-MN inhalation, with latencies increased by ∼1-2 seconds (**Figure 4C**). The ANOVA confirmed that tail withdrawal latencies were significantly affected by Time [F (3, 21) = 8.11; P<0.001], but not by Drug Treatment Condition or by the interaction of Time with Drug Treatment Condition. The post-hoc test of the Time factor confirmed that latencies were significantly elevated relative to the pre-treatment value 60 minutes after both 10 and 30 mg/mL 6-MN concentrations and 35 minutes after initiating 30 mg/mL inhalation.

### 3.5 Experiment 5: Self-administration of 6-methyl nicotine by vapor inhalation

Mean vapor deliveries obtained during self-administration did not differ between groups in the last nicotine session prior to starting the present studies, with the group assigned first to Nicotine responding for 7.8 (SEM 1.44) vape deliveries and the 6-MN group 8.4 (SEM 1.98) deliveries.

However, in the first seven experimental sessions for this study, the two groups diverged. There was a significant difference in vape deliveries associated with effect of the interaction of Group with Session [F (3.407, 61.32) = 2.85; P<0.05] in the first seven sessions and a significant effect of Group [F (1, 18) = 4.65; P<0.05] on vape deliveries in the second seven sessions (**Figure 5A**). Analysis of the percentage of responses on the drug-associated lever confirmed a significant effect of Group [F (1, 18) = 12.53; P<0.005] in the sessions 8-14, but no significant differences in the first seven sessions. Analysis of the 7 session averages (**Figure 5 B,D**) confirmed a significant impact of Group [F (1, 18) = 4.74; P<0.05] on vapor deliveries and a significant impact of Group [F (1, 18) = 9.03; P<0.01] and the interaction of Group with Session [F (1, 18) = 5.26; P<0.05] on the percentage of drug-associated responses.

**Figure 5:**
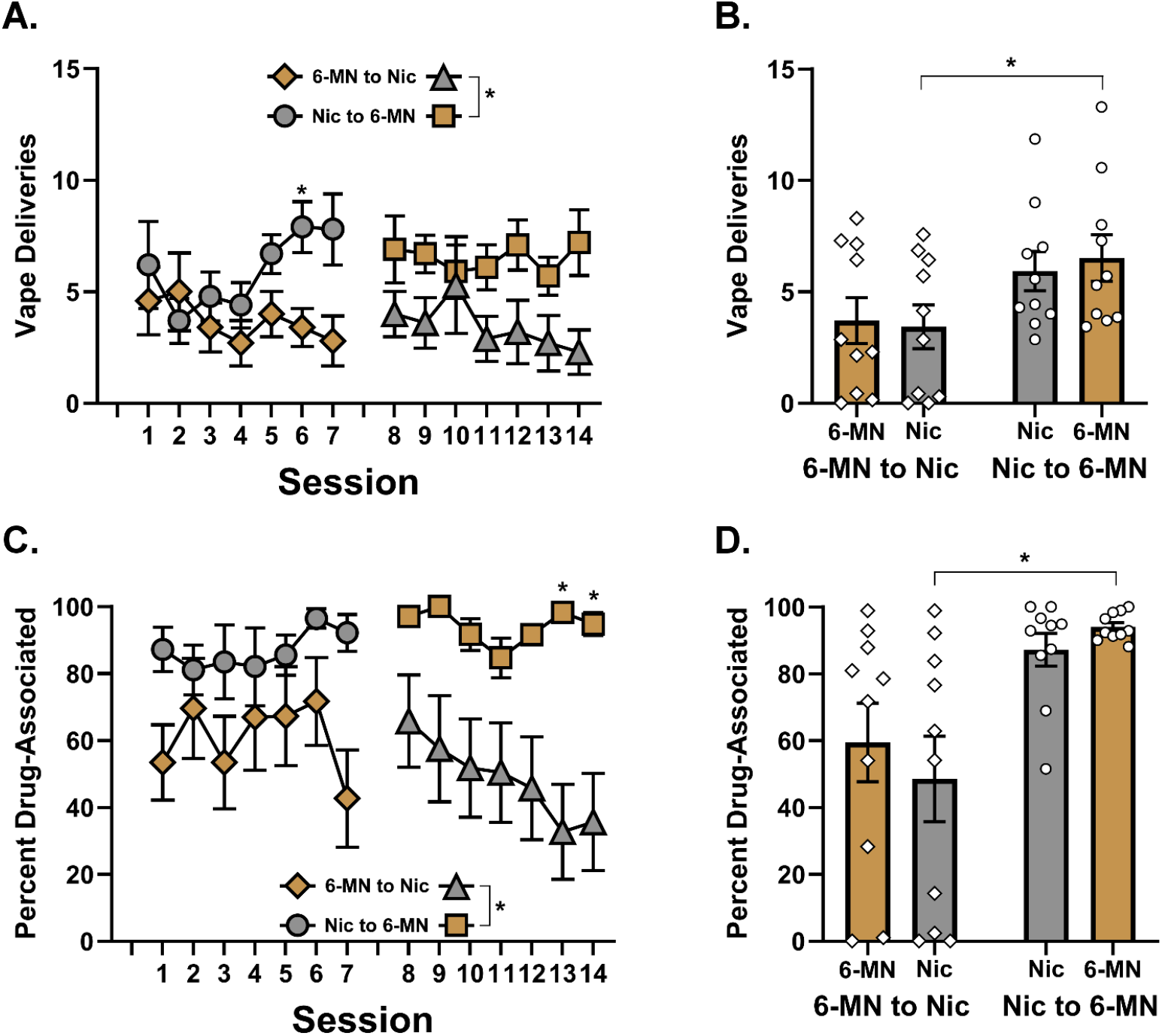
Mean vapor deliveries (**A**,**B**) and percent of responses directed to the drug associated lever (**C**,**D**) for groups responding initially for nicotine (N=10) or 6-MN (N=10) for 7 sessions and then switched to the alternate drug. Data are presented by Session (**A**,**C**) and as a 7 session average with individuals depicted (**B**,**D**). A significant difference between groups is indicated with *.

Follow-up analysis of the seven sessions for each drug within each Group did not confirm any significant difference in vape deliveries associated with the drug identity, although there was a significant effect of session [F (2.967, 26.70) = 3.03; P<0.05] confirmed for the group that self-administered nicotine first.

## 4. Discussion

The study found that 6-methyl nicotine (6-MN) administered by subcutaneous injection produced antinociceptive, thermoregulatory and suppressive effects on spontaneous wheel activity that are similar to those produced by nicotine, when injected in female rats. In addition, exposure to 6-MN by vapor inhalation also reduced body temperature and increased tail withdrawal latency to an extent similar to effects reported previously for the vapor inhalation of nicotine. Finally, this investigation showed that 6-MN substituted for nicotine in vapor self-administration, resulting in identical behavior at the same concentration. Overall, these data suggest that the behavioral effects of 6-MN are likely to be highly similar to those of nicotine.

It has been previously shown that vapor inhalation of nicotine (30 mg/mL) reduces the temperature of male and female rats (Javadi-Paydar et al., 2019; Javadi-Paydar et al., 2024), as does nicotine injection by i.p. (Dilsaver and Davidson, 1987) and s.c. routes (Javadi-Paydar et al., 2019; Javadi-Paydar et al., 2024). Here we show that these effects are also produced by 6-MN with an approximately equal effect after a 0.8 mg/kg, s.c. dose of either racemic 6-MN or (-)-nicotine in female middle-aged rats (**Figure 1**). The inhalation of vapor produced by 5-30 mg/mL of (-)-6-MN for 30 minutes reduced body temperature in a concentration-dependent manner (**Figure 4A**) with a maximum mean effect of ∼1.5°C that was similar to the effect of the injection of 0.8 mg/kg (+/-)-6-MN, s.c. (**Figure 1B**). The magnitude of the body temperature response (∼1.5°C) is also quite similar to that produced by either 30 min inhalation or 0.8 mg/kg, s.c., injection of (-)-nicotine in our prior studies in male, young adult, treatment naive Sprague-Dawley rats (Javadi-Paydar et al., 2019).

Nicotine (0.8 mg/kg, s.c.) significantly reduced the wheel activity of male and female rats in groups exposed as adolescents to repeated vapor inhalation of PG or nicotine (30 mg/mL) when assessed from PND 165-180 as young adults (Gutierrez et al., 2024b). Likewise, inhalation of nicotine (30 mg/mL) reduced wheel activity in female rats after adolescent exposure to repeated vapor inhalation of PG or nicotine (60 mg/mL) when assessed as young adults from PND 114-onward (Gutierrez et al., 2024a). The present study showed that suppression of wheel activity after injection of 0.8 mg/kg (+/-)-6-MN, s.c. matched that produced by the 0.8 mg/kg, s.c., injection of (-)-nicotine (**Figure 2**). The intermediate effect of 0.4 mg/kg (+/-)-6-MN, s.c. is consistent with the dose-dependency of (-)-nicotine for this assay (Gutierrez et al., 2024b).

Modest thermal analgesia is produced by nicotine injection when measured with tail withdrawal or hotplate response latency, however tolerance may develop with repeated nicotine exposure (Epstein et al., 1989; Yang et al., 1992). The rats in this study exhibited increased tail-withdrawal latencies after injection of 0.4 or 0.8 mg/kg (+/-)-6-MN, but not after 0.8 mg/kg (-)-nicotine (**Figure 3**). While there was no significant effect of (-)-6-MN inhalation in that group, there was a significant increase in tail-withdrawal latency in another subset of animals with no recent nicotine /6-MN exposure produced (-)-6-MN inhalation (**Figure 4C**). In this latter case, the impact was concentration-dependent, with no effects observed after 5 mg/mL inhalation, effects only 60 minutes after the start of inhalation after 10 mg/mL and at both timepoints after 30 mg/mL inhalation.

The middle aged female rats used in this study had extensive prior exposure to nicotine via early adult non-contingent exposure and ongoing self-administration of nicotine vapor. This history was considered to be an advantage as a translational model of a life-long nicotine user who decides to try the recently emerging 6-MN products. It appears obvious that the relatively recent emergence of 6-MN in products predicts that some of the most frequent use will be by individuals who have considerable prior experience smoking or vaping nicotine. There is an additional advantage to these studies in that the self-administration of drugs via any route of administration has only rarely been extended to middle age in rat models. Despite the prior nicotine exposure, the acute effects of nicotine and 6-MN were produced in these rats at relatively moderate injection doses and vapor concentrations, quite similar to those used to produce similar effects in more drug naïve rats in prior studies (Javadi-Paydar et al., 2019). The self-administration comparison also suggested an equivalency between the subjective properties of (-)-6-MN and (-)-nicotine, since each group obtained an identical number of vapor deliveries of each drug (**Figure 5**).

Analysis of ∼6 mg of 6-MN per gram of e-liquid in several products that were labeled as containing 50 mg/g 6-MN (Erythropel et al., 2024), and prior work suggest potency of 6-MN that ranges from similar to that of nicotine to 3-6 fold more potent, depending on the assay (Dukat et al., 2002; Levy and Dunn, 1979; Wang et al., 1998). Although not designed as an extensive test of relative potency, the present results do not confirm any large differences in potency in a rat model, including for physiological, reflexive and self-administration endpoints. Marketed e-liquid products claiming to be “5% strength”, or 50 mg/mL, were analyzed to contain ∼6 mg of 6-MN per gram of e-liquid which suggests a market-driven conclusion that 6-MN is much more potent than nicotine (Erythropel et al., 2024). In an *ex-vivo* assay, (-)-6-MN was about 3-fold more potent than (-)-nicotine in a competition binding assay in rat membrane (Wang et al., 1998), and *in vivo* (+/-)-6-MN was found to be about 3-fold more potent than (+/-)-nicotine in tail-withdrawal anti-nociception and 5-fold more potent in inhibiting spontaneous activity in mice (Dukat et al., 2002). Overall, however, the present studies do not support a conclusion that 6-MN is substantially more potent than nicotine after inhalation or subcutaneous injection.

There are a number of caveats that caution against overinterpretation of dose comparisons in this study. The salt form of the drug available for injection use was the racemic mixture, thus comparison with the (-)-nicotine injection data requires caution. The stereoisomers of nicotine are metabolized at equivalent rates in female rats (Jenner et al., 1971), thus differential metabolism is unlikely to explain any observed similarities or differences between racemate and the more-active stereoisomer. However, while the (+)-nicotine isomer was reported to be >100 fold less potent than (-)-nicotine when administered intraventricularly (Abood et al., 1978), (+)-nicotine was only 9 times less potent when injected s.c. in a drug-discrimination assay (Meltzer et al., 1980) and the two isomers were equipotent in evoked responses of neurons in frog olfactory epithelium (Thurauf et al., 1995). The relative potency of stereoisomers on physiological endpoints is not always as would be predicted from *in vitro* assays, for example S(+)-MDMA and the racemic produce equivalent hyperthermic effects in monkeys and mice (Fantegrossi et al., 2003; TAFFE et al., 2006). Nevertheless, while we cannot rule out the possibility that (-)-6-MN might be more potent when injected, the present studies show (-)-6-MN and (-)-nicotine produce similar effects when inhaled at the same concentrations. As a final caveat, male rats were not included in the original study, and thus were not available for this work, since we had previously found they self-administered nicotine vapor at lower rates than did females and there was minimal effect of repeated adolescent exposure to Nicotine (30 mg/mL) vapor on self-administration in males (Gutierrez et al., 2024b). Consequently, the present results may not generalize to male animals.

In summary, this study confirms that 6-MN produces effects that are highly similar to nicotine *in vivo* in laboratory rats, with comparable effects produced by comparable dosing. Effects observed after either s.c. injection or vapor inhalation accord with prior reports for nicotine administered by each route in terms of the magnitude of effect, the time-course and the approximate dose range. Thus, it is likely that the 6-MN analog used in novel consumer products will entail similar risks to health as do the nicotine-containing versions.

## Acknowledgements

The authors thank Wayne Mascarella, Ph.D. for assistance with sourcing and verifying the 6-methyl nicotine used in these investigations. These studies were supported by the Tobacco Related Disease Research Program (T33IR6653 MAT) and by the U.S. National Institutes of Health (DA057423 MAT). The funding entities had no influence on the study design, data interpretation, manuscript creation or in the decision of when and what to publish from the studies conducted.

